# On the Rotational Structure in Neural Data

**DOI:** 10.1101/2023.09.11.557230

**Authors:** Ekaterina Kuzmina, Dmitrii Kriukov, Mikhail Lebedev

## Abstract

Spatiotemporal properties of the activity of neuronal populations in cortical motor areas have been the subject of many experimental and theoretical investigations, which generated numerous interpretations regarding the mechanisms of preparing and executing limb movements. Two competing models, namely representational and dynamical models, strive to explain the temporal course of neuronal activity and its relationship to different parameters of movements. One proposed dynamical model employs the jPCA method, a dimensionality reduction technique, to holistically characterize oscillatory activity in a population of neurons by maximizing rotational dynamics that are present in the data. Different interpretations have been proposed for the rotational dynamics revealed with jPCA approach in various brain areas. Yet, the nature of such dynamics remains poorly understood. Here we conducted a comprehensive analysis of several neuronal-population datasets. We found that rotational dynamics were consistently accounted for by a travelling wave pattern. To quantify the rotation strength, we developed a complex-valued measure termed the gyration number. Additionally, we identified the parameters influencing the extent of rotation in the data. Overall, our findings suggest that rotational dynamics and travelling waves are the same phenomena, which requires reevaluation of the previous interpretations where they were considered as separate entities.

---

In motor cortical areas, such as the primary motor cortex (M1) premotor cortex (PMC) and supplementary motor area (SMA), most neurons modulate their firing rate in association with preparation and execution of limb movements.

Nowadays, neurophysiologists strive to record from large populations of single neurons, which allows to better understand their ensemble properties. By varying motor tasks and recording from different subdivisions of the brain motor network, one can assess the composition of population activity and the information it encodes and processed.

Numerous experimental and theoretical have tackled the function of cortical motor networks, and several models and interpretations have been proposed. The representational model that dates back to the time when neurophysiologists recorded from one neuron at a time posits that neuronal firing rate in motor areas represent various parameters, such as spatial target location, limb kinematics, and muscle force [1–5]. The representational model has been widely used to interpret neurophysiological results and it has been practically tested in brain-computer interfaces (BCIs) that converted neural activity into the parameters of prosthetic limb movements [6–9].

Yet, the representational model cannot account for the considerable variability of neuronal activity in the motor cortex. Neural representation can change across movements, contexts, and behaviors [10]. Particularly, temporal patterns of single-neuron activity considerably vary whose directional tuning patterns could be very different during preparation versus execution of limb movements [11]. To account for these peculiarities, more parameters should be explicitly added to the representational model – a reductionist approach that raises questions about the model’s reliability [10].

An alternative interpretation of the highly complex activity of neurons in the motor and premotor cortex is based on the dynamical-system approaches where an emphasis is put on the collective activity of neuronal populations. Thus, the approach called optimal feedback control (OFC) describes how we move and utilize peripheral feedback without explicitly specifying each neuron’s activity in terms of represented motor variables [12–17].

In 2012, Churchland and colleagues developed a model that, like OFC, describes neural population activity using dynamical systems techniques but omits the sensory feedback term. In its simplest form, the model can be expressed as: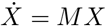, where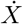 represents the time derivative of the neuronal state space vector, *M* is the dynamical system matrix, and *X* is the neural state space vector.

The dynamical model suggests that neurons in the motor and premotor cortex do not so much represent movement parameters as keep an action going [18]. Neurons become active when movement is prepared [17], with the subsequent dynamics solely governing movement progression.

By constraining *M* to possess a skew-symmetric structure, the dynamical system yields rhythmic, oscillatory temporal patterns, known as rotational dynamics. Churchland et al. discovered these patterns in the motor cortex population activity of rhesus monkeys using joint Principal Component Analysis (jPCA), a linear dimensionality reduction method that projects data to the space maximizing rotational pattern and captures temporal dependencies between latent states [19]. The projections of arm-reaching data were found to resemble curved trajectories in state space, indicative of a dynamical system that provides a “basis set for generating the necessary patterns of muscle activity” [20].

Since the original publication of Churchland and his colleagues, the rotational dynamics perspective and the jPCA method have gained a widespread popularity in hundreds of subsequent publications. Rotations of low-dimensional jPCA (or PCA) projections have been studied across various brain areas and species (Fig. 1).

**Figure 1:**
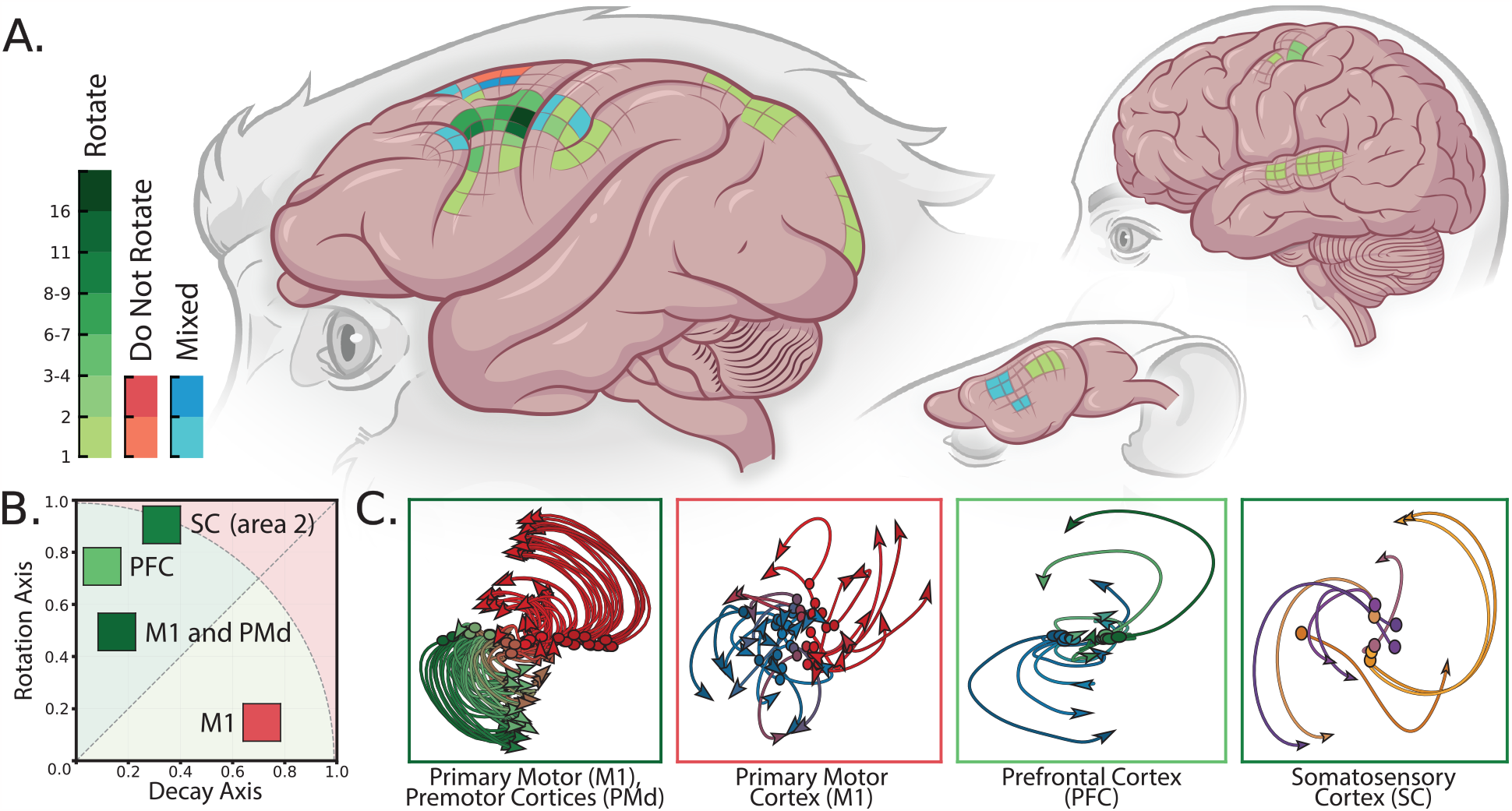
A) The brain areas that were explicitly studied with rotational dynamics approach in rhesus monkeys, humans and rodents. Green or red colors indicate the discovered presence or absence of rotations respectively, while blue color depicts contradictory results. The intensity of the colors correspond to the number of papers examining this area of the brain. B) The complex-valued measure showing the extent of rotational dynamics forms the gyration plane. C) Examples of jPCA projections for different brain areas from datasets that were used in our study. The details of the datasets can be found in Supplementary Fig. S3

In monkeys, the following brain areas have been studied using jPCA: primary motor cortex (M1) and premotor cortex (PM) [20–38]; somatosensory cortex (S1) [22, 24–26]; supplementary motor area (SMA) [28, 29, 39, 40]; primary visual cortex (area V1) [21]; frontal cortex [39, 40], prefrontal cortex [41–43]; and posterior parietal cortex [44]. In humans, the following areas have been studied: primary motor cortex (‘hand knob’ area) [45, 46] and left superior temporal gyrus (STG) [47] in individuals with movement impairment. In rodents, jPCA method has been applied to the data collected in the following areas: auditory and motor cortices of mice [48–50] and primary motor cortex and dorsolateral striatum (DLS) of rats [51].

Furthermore, jPCA method has been applied across different motor tasks and contexts. In monkeys, the following tasks have been studies: center-out hand-reaching movements [20–22, 25, 26, 32, 52] (cue-initiated and self-initiated [27, 29, 35], executed and observed [32], instructed-delayed [30, 44], with corrective movements [36]); reach-to-grasp hand movements [31, 33, 35, 53]; isolated [22] and delayed grasping movements [37, 38]; reach-grasp-manipulate movements [23, 34]; and cycling hand movements [28]. Additional tasks include posture perturbation [24], time interval estimation [39, 40], decision-making using working memory [41], perceptual decision-making [42, 43], perception of cyclic visual stimuli [21], and neural activity during sleep and sedation [35]. In individuals with movement impairment, the following paradigms have been used: finger-reaching movements [45], speech and orofacial movements [46], and listening to speech [47]. In rodents, the following tasks have been studied: mice during reach-to-grasp movement [50], listening to auditory stimuli [48] and decision-making tasks [49]; rats during reach-to-grasp movements [51].

Despite the growing body of work exploring neural rotational dynamics across different brain areas, species, contexts, and movements, noticeable inconsistencies in the results and interpretations have emerged.

Although various metrics exist to measure rotations in data [20, 22, 35, 40, 41], a precise definition of rotational dynamics remains elusive. In the original paper, rotations were defined as “consistent ordered state-space rotations across different reaching movements” [20]. Some authors considered trajectories rotational if specific requirements were met [42, 45], but often, studies claimed the presence of rotational structure based on visual evaluation or *R*^2^ fitting coefficients [24, 29, 32, 34, 35, 37, 44, 47]. Visual estimation of rotational dynamics, combined with limited recorded movement conditions [22, 25, 26, 45, 46], could yield ambiguous results.

Furthermore, no consensus exists on the causes of rotational dynamics and the significance of state-space rotations. Initially, rotational dynamics implied an intrinsic, autonomous dynamical system [17, 18, 20], which can be disrupted by movements, requiring more inputs from other brain areas [22]. However, subsequent work proposed alternative interpretations: the rotational dynamics represent the opposite - online control of motions [24], sensorimotor feedback control and/or a cognitive strategy [35, 36, 54].

Interpretation difficulties extend beyond the motor cortex. For example, the supplementary motor area (SMA) neurons exhibited varying rotational patterns depending on the task. SMA neural activity did not exhibit rotations during reaching tasks [29], had rotational “helical population trajectory” during cycling hand movements [28] and rotated during time interval estimation task [40].

It was suggested that the absence of rotational dynamics in SMA reflected the fact that “SMA keeps track of context and must robustly differentiate between actions that diverge with time” [55]. However, the same interpretation was used to explain why rotational dynamics were present in the superior temporal gyrus (STG): “structured rotational dynamics can act as clocks” and “indicate transition from one stimulus to the other” [47].

Many studies argued that rotational dynamics were generated due to the intrinsic recurrent connectivity of M1 and PMd [18, 22, 39, 48, 55, 56], with recurrent neural networks (RNN) often used to validate this hypothesis. While RNNs (trained to produce muscle activity from neural activity) generated emergent rotational dynamics [18, 41, 57–59], one study found that “rotational dynamics were generated in networks trained with and without intrinsic recurrent connections”, which questions the connectivity hypothesis [24].

Some works suggested that rotational trajectories could be artifacts of the jPCA method. Churchland et al. proposed that “trivial rotations could emerge due to the method’s power in finding state-space rotations for diverse and multiphasic responses” [20]. To validate rotation importance, they employed data shuffling procedures, rearranging neurons between conditions.

Additionally, Michaels et al. found that jPCA revealed trivial rotations for the representational view model, and the original shuffling procedures failed to differentiate between them [18]. They proposed a more complex shuffling procedure, preserving the neuron covariance matrix while rearranging neurons between conditions.

Subsequently, it was observed that previous shuffling procedures destroyed data rather than identifying triviality, leading to an even more complex shuffling method confirming that dynamical structure was not a byproduct of simpler features [60].

Several publications questioned that state-space rotations describe any meaningful physiological functions and instead represent a visualization tool whose interpretation remains obscure without additional analyses. Thus, Lebedev et al. demonstrated that rotations in neural population activity could represent spatiotemporal neural patterns best described as travelling waves [61]. Travelling waves are frequently encountered in neural datasets [62–66], including motor learning data [67], and reach-to-grasp movements [51, 68], as well as datasets for RNNs trained for muscle activity production [69] and decision-making [41]. Additionally, Proix et al. [70] demonstrated that a temporal covariance matrix structure creates a *horseshoe artefact* - a frequently encountered consequence of dimensionality reduction methods that yields rotations of state-space trajectories.

Despite these ambiguities, rotational phenomena appear in many brain areas, movements, contexts, and behaviors. This ubiquity raises questions about the commonalities between all neural population activities mentioned.

Most existing works explore rotations heuristically. In contrast, our approach is purely data-driven: abstracting from assumptions about the data, like underlying dynamical or representation view models; and freeing from any interpretation of rotational dynamics, like an “engine of movement” [20] or a “spring box” [18]. In our work, we seek an answer to the question: “Why does neural data rotate?” or, more formally, “Which neural data properties are *necessary* and *sufficient* for rotations to occur?”

In order to answer this question, we analyzed the phenomenon of rotations, explaining the *necessary* and *sufficient* requirements for its occurrence. We distinguished between individual rotations of one condition and structural rotations of all conditions together, and identified the sources of rotational dynamics. We introduced the gyration number, a complex-valued measure, to study neural datasets and detect structural rotations, and enable comparison between datasets. We used a simple structural rotation model to examine the gyration number properties and identify the data characteristics that influenced the pattern of structural rotation.

## 1. Results

We began by distinguishing between two concepts: *condition-dependent rotation* and *structural rotation*. The former referred to a single condition’s rotation resulting from the neural dynamics of that condition, while the latter denoted the tendency of multiple conditions to form a co-directional rotational pattern (similar to a “sheaf” of trajectories). It was crucial not to confuse these two concepts of ‘rotations.’ We separately examined conditional and structural rotations, uncovering their nature and providing the necessary and sufficient conditions for their occurrence. Most importantly, we highlighted the limitations of dimensionality reduction methods in analyzing neural population dynamics data (Fig. 2A).

**Figure 2:**
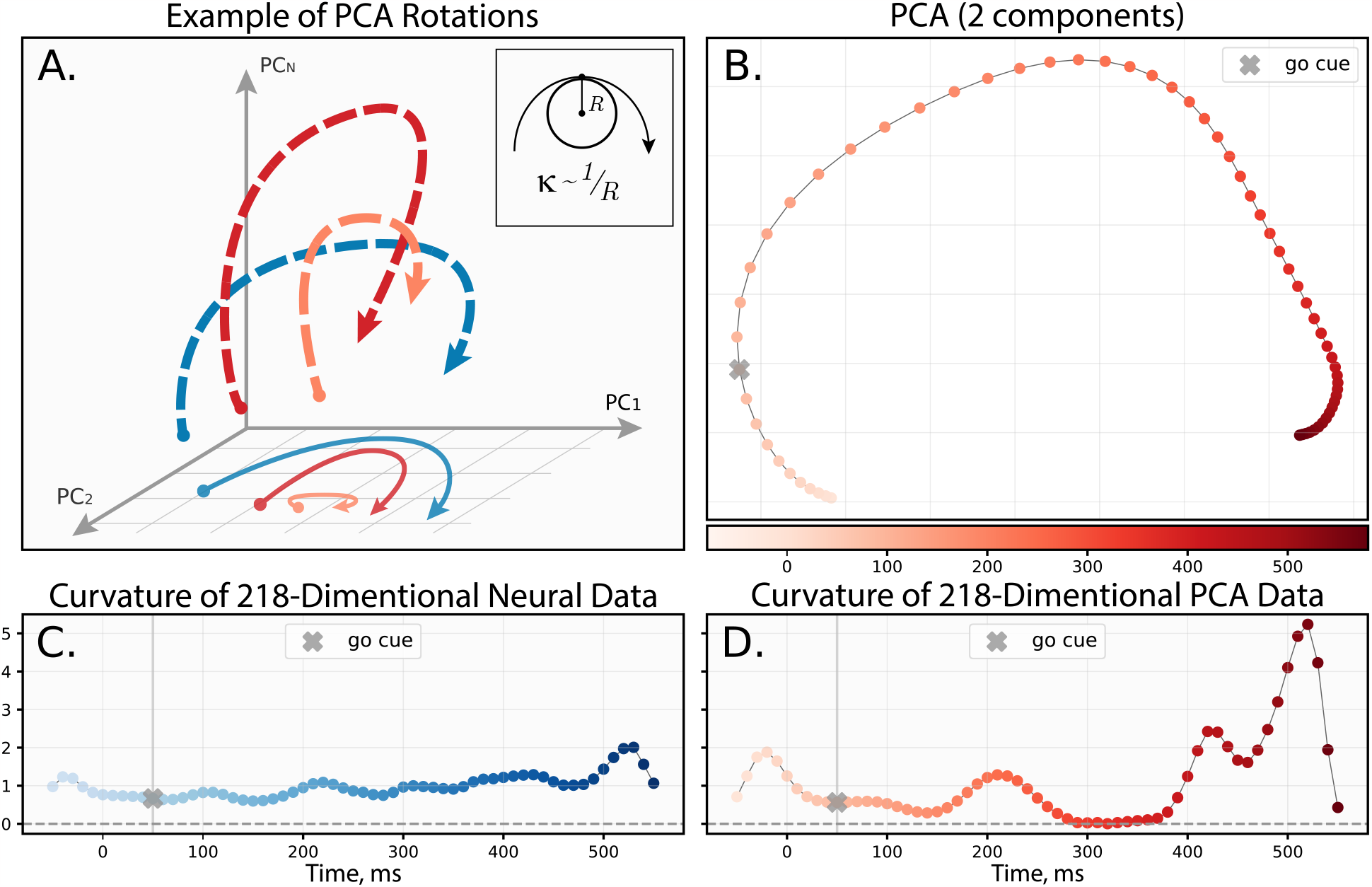
The necessary requirements for one condition rotation. A) A schematic illustration of how dimensionality reduction techniques could distort the original shape of the trajectory. The embedded plot demonstrates the concept of curvature graphically. B) Projection of one condition onto the PCA1-2 space. C) The curvature of the original 218-dimensional curve changing across time. D) The curvature of the projected 2-dimensional curve changing across time. “Go cue” point refers to the movement onset time point. The data for panels B, C, and D are taken from [20].

### 1.1. Necessary requirements for one-condition rotation

A necessary requirement refers to a parameter estimated from the data that unequivocally signifies the presence of rotations. Any one-condition neural recordings have a form of time-dependent vector that can be also treated as a point moving in high-dimensional space and forming some kind of trajectory. To answer the question “Is the trajectory rotational?” we had to define the concept of ‘rotations’ in general and develop a method to measure them. Several studies proposed to measure an angle between the radius vector to point trajectory and its velocity vector [20, 41, 42, 45]. Which detects the ‘rotational’ patterns in general, but does not give much information about their magnitude. Instead of measuring the angle, we propose to measure the curvature of a trajectory.

Curvature is a measure of non-collinearity between velocity and acceleration vectors that has a differential measure of trajectory rotation at each point. Like angle, curvature has a clear interpretation as the inverse radius of the inscribed circle (*κ* ∼ 1*/R*_*κ*_) tangent to the given point on the trajectory (Fig. 2A, embedded image). Consider a time *t* parameterized space curve in *n* dimensions (with *n* representing the number of neurons in a condition) given in Cartesian coordinates by *x*(*t*) = [*x*_1_(*t*), …, *x*_*n*_(*t*)]^*T*^ . The original expression for curvature is as follows:

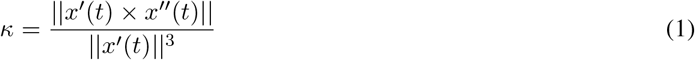

The presence of the cross product (×) in the numerator of the formula poses a challenge. It is well-known that the cross product is poorly defined for dimensions other than 3 or 7 [71]. Consequently, computing the result of the cross-product explicitly for an *n*-dimensional space is problematic. Fortunately, it is enough for us to estimate only the norm of curvature in order to understand its magnitude, which results in a significant simplification of the equation:

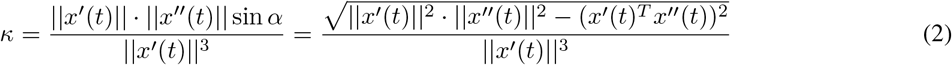

where *α* represents the angle between the velocity and acceleration vectors. Consequently, the output is the absolute value of the curvature. Although this could pose an issue when analyzing data with a change in the direction of rotation (e.g., an S-shaped trajectory), this is not an issue here because neural population data consistently exhibit strictly positive curvature (see Supplementary Figure S3).

Upon deriving the expression for the curvature, we estimate it directly from the neural population data by calculating numerical gradients along the trajectory. An example of a one-condition projection on PCA1-2 space is shown in Figure 2B, with the corresponding curvature calculated for both the curve in the original *n*-dimensional space (Fig. 2C) and the projected curve (Fig. 2D). This example illustrates how curvature can change when applying dimensionality reduction techniques. In the original *n*-dimensional space, the curve exhibits strictly positive curvature, which should be interpreted as turning in one direction. The varying curvature magnitudes represent sharper turns in high-dimensional space. It is worth noting that these turns may disappear following the projection procedure (Fig. 2A). Therefore, we argue that the curvature profile provides an insight into the actual neural dynamics than its angle-based counterpart. Furthermore, a non-zero curvature value is a necessary requirement for the one-condition rotation, as it directly indicates a non-zero angle between the acceleration and velocity of a moving point.

### 1.2. Sufficient requirements for one-condition rotation

A sufficient requirement refers to the presence of a structure in the data that generates the observed rotational dynamics.

We begin by examining the approximation of neural population dynamics proposed in [61]. By ordering neurons based on their spike-train peaks in the average PETH diagram (Fig. 3A, Suppl. Fig. S3), we observe a travelling wave pattern in the data (spreading across the neural network). In this work, the authors speculated that such a “running wave” pattern is sufficient for generating rotational dynamics and even proposed a simpler model (referred to as “naive”) consisting of a basic Gaussian function (Supplementary figure S6A). The underlying Gaussian function takes the following form:

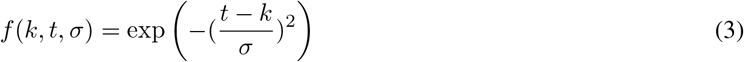

where *k* is the time point of wave peak, *σ* determines “width” of the wave. One important observation is that the parameter *k* can be expressed as a function from the ordered indices of neurons, namely:

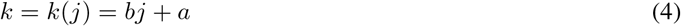

where *j* is an index of a peak-ordered neuron; *b* is a parameter determining incline of the wave on the Figure (Fig. 3A), we will also refer to it as a wave speed; *a* is an initial shift of the wave. Previously, it has been shown that if *b* = 0 the wave does not propagate and the related rotational dynamics collapses (Supplementary figure S6B, C, D) [61]. This is due to the activity of all neurons being strongly correlated and the corresponding covariance matrix (which is used for PCA computation) becoming singular. The other extreme case arises when the activity of neurons do not correlate (Supplementary figure S6E, G, F) with each other (*b* → ∞) leading to a diagonal covariance matrix and an absence of any rotational structure in the data (eigenvectors of such covariance matrix are just columns of identity matrix).

**Figure 3:**
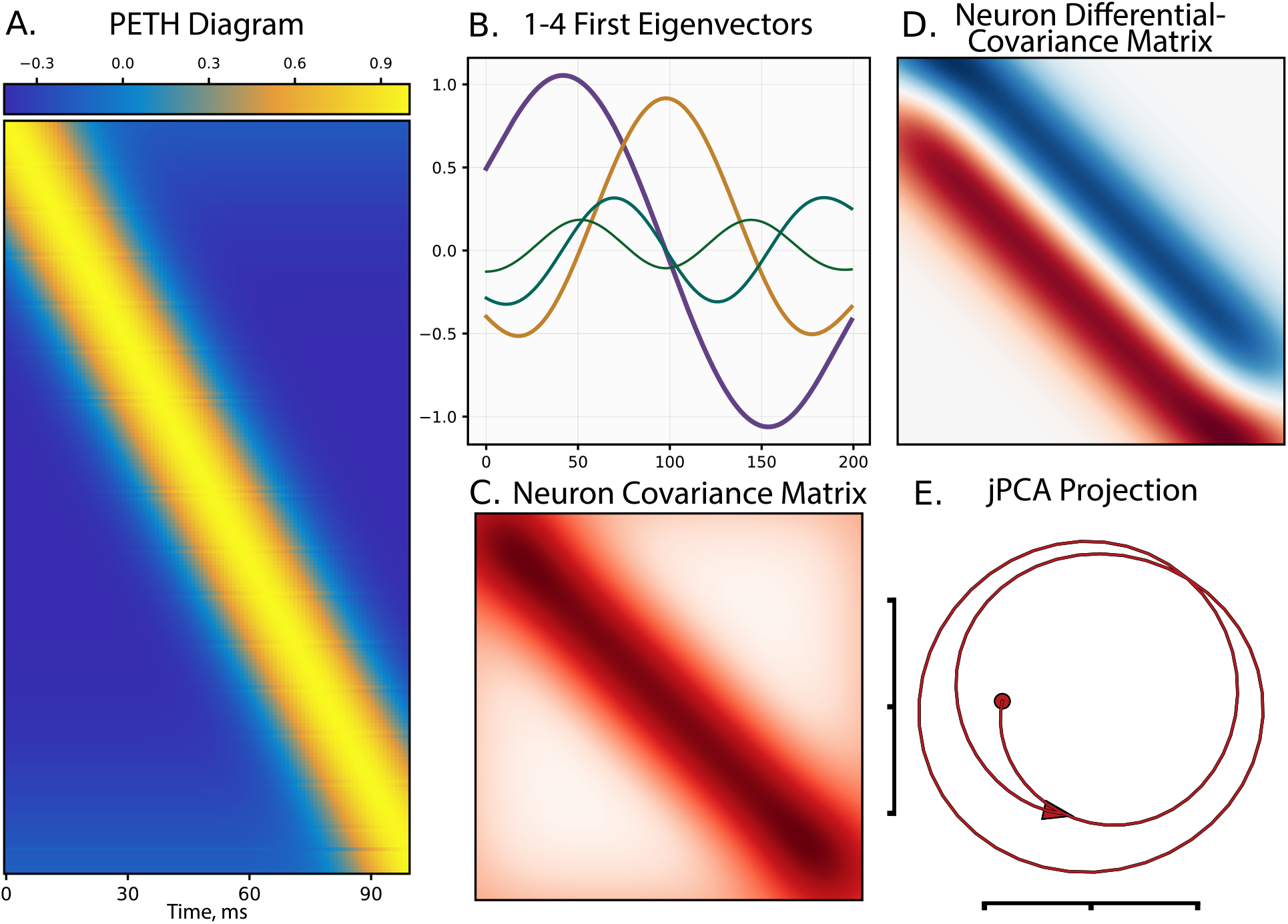
The sufficient requirements for one condition rotation. A) Peri-event time histogram (PETH) of the simulated Gaussian travelling wave. B) Oscillating eigenvectors of the Toeplitz Neuron covariance matrix structure corresponding to the simulated travelling wave. C) Toeplitz covariance matrix for the simulated travelling wave. D) Toeplitz-like differential covariance matrix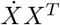 for the simulated travelling wave. Small distortions at the corners of the matrix are related to the cutoff of the wave at the edges of the PETH. E) jPCA projection of the PETH data demonstrates the proper rotational dynamics.

To elucidate the origins of rotational dynamics in the travelling wave structure, we propose two types of evidence: weak and strong, which demonstrate the sources of such dynamics. For the weak evidence, we need to examine the following extreme case in which only two neighboring (by index) neurons are correlated. In our naive model example, this case implies a tridiagonal Toeplitz feature covariance matrix (of size *n* × *n*, where *n* is the number of neurons). This matrix possesses a remarkable property: its eigenvalues and eigenvectors can be expressed in a closed form [72], specifically:

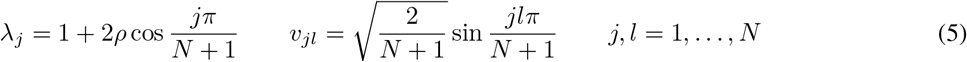

where *ρ* is the correlation coefficient between two neighboring neurons, and *l* is the index of a component of the *j*-th eigenvector. We can see from the eigenvector expression that it contains a sinusoidal function with an increasing period as the corresponding eigenvalue index increases. Plotting the first and second eigenvectors of the tridiagonal covariance matrix provides insight into the nature of rotational dynamics in naive model-derived data [73], where we observe a cyclic pattern, also known as a “horseshoe” [70], generated by two orthogonal sinusoidal vectors with different periods. Consequently, the minimal model with the simplest covariance structure (conditioned by the travelling wave) generates oscillating eigenvectors (Figure 3B), explaining the rotational structure of projected data. We can generalize this approach by adding diagonals to the feature covariance matrix, which corresponds to decreasing the *b* coefficient in Equation 4. It can be demonstrated that the Toeplitz covariance matrix (Figure 3C) also preserves the oscillating eigenstructure in this case. Moreover, this fact can be proven rigorously for the generalization of Toeplitz to Circulant matrices [74].

The strong evidence of rotational structure in travelling wave data arises from another object implicitly used in the jPCA procedure. The original paper describes jPCA as an approach to finding a projective space in which rotations are most explicit [20]. To achieve this, the authors suggest finding a system matrix in the equation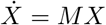 under the constraint that matrix *M* is a skew-symmetric matrix, i.e., *M* = −*M* ^*T*^ (referred to as *M*_*skew*_). This property implies that all eigenvalues of this matrix are purely imaginary, meaning that matrix *M*_*skew*_ acts as a rotation operator on *X* without inflating or contracting the corresponding trajectory. To confirm this fact, we can consider the solution to the jPCA problem in continuous time:

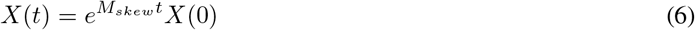

It is known that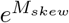 is an orthogonal matrix and, therefore, does not inflate (or contract) the trajectory *X*(*t*) over time. Thus, if neural population data have a specific structure implying a skew-symmetric system matrix, the data automatically generates rotational dynamics. What structure should the data have to imply a skew-symmetric *M* ? We demonstrate that it is enough to examine the structure of the so-called *time differential covariance matrix* 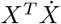 [75].

**Theorem 1**. *Let X* ∈ ℝ^*n×t*^ *and M* ∈ ℝ^*n×n*^. *Then, if*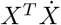 *has a skew-symmetric structure, then M is skew-symmetric*.

*Proof*. Let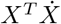 is skew-symmetric and 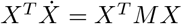, then

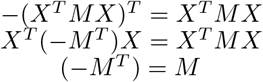

Figure 3D demonstrates that travelling wave data generated from our model (eq. 3) yield almost (omitting edge effects) skew-symmetric neuron differential covariance matrix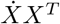 (the corresponding time differential covariance matrix is depicted in Supplementary Figure S5). Moreover, we can show rigorously that the model 3 generates skew-symmetric neuron differential covariance matrix satisfying the definition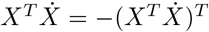 (see supplementary derivation A). Thus, the generated data implies rotational dynamics on the jPCA plot (Fig. 3E).

Thus, the proposed weak and strong evidence comprehensively demonstrate the presence of rotational dynamics in the data with a travelling wave structure. Therefore, we contend that the presence of a travelling wave in a neural population data is a sufficient requirement for one-condition rotation but not for structural rotations discussed below.

### 1.3. Requirements for structural rotations

To better understand structural rotations, we constructed a complex-valued measure that can detect structural rotations in any time-course data, irrespective of the underlying model. We will then examine how the naive travelling wave model can generate such data and identify key parameters of this model that influence the occurrence of structural rotations.

#### 1.3.1. Model-free approach

Distinct experimental conditions produce varying neural population behaviors. However, when one condition is similar to another, they exhibit similar dynamics. When considering these similar conditions collectively, we can envision them as a “sheaf” of trajectories in a multidimensional space. Dimensionality reduction techniques, such as PCA or jPCA, enable the visualization of multidimensional trajectories by projecting the original data onto a 2 or 3-dimensional subspace spanned by vectors that maximize variance (PCA) or the rotational component (jPCA). The standard jPCA procedure involves stacking different conditions within a single tall matrix in order to perform dimensionality reduction and then unstacking to reveal the desired “sheaf” pattern [20]. However, the “sheaf” pattern is usually neglected due to the high similarity across the conditions. Consequently, the cross-conditional mean is usually subtracted to eliminate similarities and highlight differences between conditions, which is nearly equivalent to subtracting the first principal component. This is where structural rotations could emerge. High correlation of conditions and sufficient variance between conditional-mean values (Suppl. Fig. S1) results in the formation of the low-dimensional manifold that exhibits rotations which could be detected by jPCA. However, in a classic jPCA problem, the matrix *M* fits the PCA-compressed data losing a part of the information. Otherwise, when fitting to full data, the method may experience convergence problems.

Next, we consider the question whether it is it possible to capture the low-dimensional rotation pattern with a single-measure calculation from the complete dataset. In the previous paragraph, we provided a preliminary consideration of this question by examining the time differential covariance matrix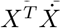, which is directly related to the form of matrix *M* . It can be shown that if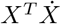 exhibits skew-symmetry, then,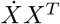 will also possess this property (Suppl. Fig. S5) -we refer to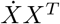 as the spatial differential covariance matrix. This matrix has a useful property: its eigenspectrum can detect low-dimensional rotations. Specifically, we can propose the following empirical rule: if the first pair of complex-conjugated eigenvalues predominates compared to the power of the entire spectrum, and the imaginary parts of these eigenvalues are greater or approximately equal to their real parts, the entire dataset will exhibit structural rotations, i.e.

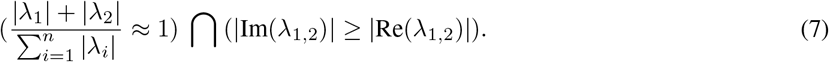

Thus, the data displaying a purely skew-symmetric spatial differential covariance matrix exhibit “ideal” structural rotations (which we discuss below). In the experimental datasets, the matrix is not entirely skew-symmetric and has a symmetric component that affects the corresponding real part of eigenvalues. When the eigenvalues possess a small negative real part (relative to the imaginary part), the jPCA of the data reveals a spiral-like decaying pattern. In contrast, a positive real part corresponds to an expanding spiral. However, if the real part of the eigenvalues becomes too large, the linear projective methods capture something other than a rotational pattern, resulting in the destruction of the observed rotation. In the limiting case, the differential covariance matrix may have purely real eigenvalues - in such a scenario, no rotational structure can be observed.

In light of these considerations, it is natural to allocate neural population dynamics datasets in the space spanned by real and imaginary parts of their first pair of complex-conjugated eigenvalues normalized by the spectrum power (Fig. 5) - we will refer to this approach as *Gyration Plane*. The normalized imaginary part measures the structural rotation component in the data and the normalized real part measures the inflation/deflation component of the data. This visualization approach provides a holistic view of the rotational dynamic problem allowing a comparison of different datasets with each other in the model-agnostic (no fitting as in jPCA) manner not requiring a dimensionality reduction procedure.

#### 1.3.2. Model-based validation of the approach

The considerations above can be tested using the naive model of structural rotations - the travelling wave model. We have already described this model for single-condition rotation, and it can be easily generalized to produce structural rotations. The primary motivation for this generalization is to explore different parameters of this model to better understand dataset behavior in the Gyration Plane. Let *D* ∈ ℝ^*c×t×n*^ represent a dataset of neural population dynamics across *c* conditions. Based on relations 3 and 4, we can formulate the following model to generate the dataset:

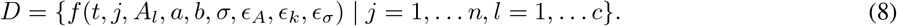

The function *f* is an extended version of the relation 3, specifically:

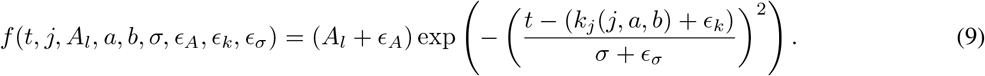

Here, six parameters of the naive model define a specific synthetic dataset. *A*_*l*_ represents the average amplitude of a wave in a given *l* condition, which can be perturbed with random noise *ϵ*_*A*_ ∼ ℕ (0, *σ*_*A*_) within the condition. *k*_*j*_ = *bj* +*a* is defined as the phase (or shift) of the travelling wave, depending on neuron index *j*, wave speed *b*, and initial phase *a*; this can now be perturbed with phase noise *ϵ*_*k*_ ∼ ℕ (0, *σ*_*k*_). *σ* is the average “width” of the wave in a specific condition, which can be perturbed with random noise *ϵ*_*A*_ ∼ 𝕌 (0, *σ*_*σ*_). Overall, this model represents a set of *c* travelling waves with a particular average amplitude *A*_*l*_ that varies across conditions.

Figure 5B displays multiple datasets *D* generated by various parameter sets. Each embedded miniature represents the jPCA visualization of the corresponding dataset, positioned according to its coordinates in Rotation Space. The coordinates can be computed using a three-step algorithm: (i) stack the conditions of matrix *D* into the tall matrix *X* ∈ ℝ^*c·t×n*^; (ii) identify the eigenvalues of the squared matrix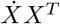 and (iii) calculate the coordinates *x* and *y* in Rotation Space using the following formulae:

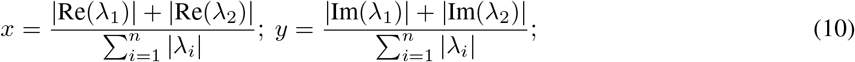

where *λ*_1_ and *λ*_2_ are the first pair of complex conjugated eigenvalues (or the first two largest eigenvalues for the case of real eigenspectrum).

To investigate how varying model parameters affect the position of a dataset *D* in the gyration plane, we begin with noise-free synthetic datasets comprising 8 conditions (chosen as it is the most common number of conditions found in the literature [18, 22, 24–26]), 200 neurons (*n* = 200), and *A*_*l*_ ranging from 0.5 to 1.0. All preprocessing steps for jPCA construction, including cross-conditional mean subtraction and soft-normalization, are maintained. Miniatures with green borders display noise-free cases where only wave speed *b* and wave width *σ* parameters are altered. We observe that decreasing travelling wave speed transforms structural rotations toward a spiral-like pattern results in collapsed rotations as wave speed approaches zero. Conversely, simultaneously increasing wave speed and decreasing wave width results in pure imaginary rotations (leftmost green miniature), corresponding to a purely skew-symmetric spatial differential covariance matrix. This leads to our first conclusion: the imaginary part of the first eigenvalue is associated with the roundness of the dataset, whereas the real part is related to its stretchiness.

Introducing noise shifts the dataset towards the origin of the gyration plane and distorts the trajectories, which leads to the second conclusion: adding noise increases the dataset’s intrinsic dimensionality. Indeed, a noise-free dataset can be well-described by two complex-conjugated jPCA components capturing pure rotations (round or stretched). In contrast, noise-contaminated travelling waves necessitate more components for accurate description. The rotation space captures noise-affected datasets in the vicinity of the origin, where the magnitudes of the first pair of complex-conjugated eigenvalues are no longer comparable with the entire spectrum (condition 7).

We tested different numbers of conditions (Fig. S7A) and found that the relationships between datasets are preserved. We also tested different numbers of neurons (Fig. S7C), with relationships between datasets remaining consistent. Altering the range of condition average amplitudes (*A*_*l*_) makes the entire dataset more resistant to noise (Fig. S7B). A change in the wave shift reduces the decay component (Fig. S7D). Thus, through the series of such experiments, a continuous change occurs in gyration number under the continuous change in the travelling wave parameters.

#### 1.3.3. Testing the approach on experimental datasets

We compared five experimental datasets containing 59 recordings in total, using the gyration number measure (Fig. S4 and Fig. 4). These datasets varied in sampling frequency, movement timings, number of conditions, and neurons, and include diverse brain areas such as motor and premotor cortex (M1 and PMd), somatosensory (S1), supplementary motor area (SMA), prefrontal cortex (PFC), and parietal cortices. More detailed description of the datasets can be seen in Supplementary Table 1. Four of the datasets were recorded from rhesus monkeys during hand-reaching movements while one dataset contained the recordings from SMA during grasping movements, where presumably the dynamics were non-rotational. We preprocessed the data according to the source papers and allocated all datasets in the gyration plane.

**Figure 4:**
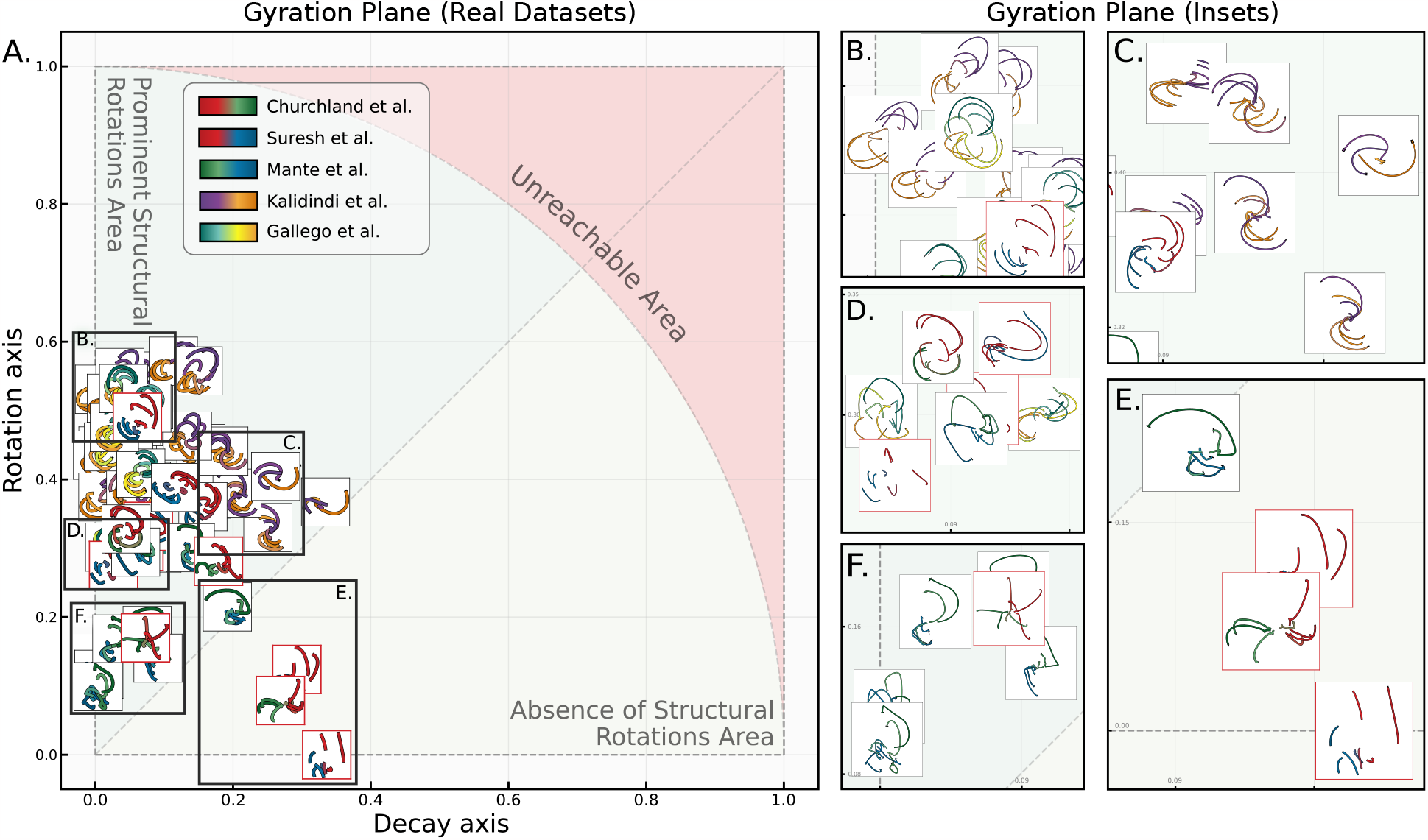
The gyration plane for real neural population dynamics datasets. A) Gyration plane - the visual representation of the developed complex-valued measure of structural rotations - gyration number. The real part of the gyration number corresponding to the decay axis shows the strength of the “stretching” force in the dataset which can be negative - trajectories contract, or positive - trajectories inflate. Both types deteriorate a pattern of structural rotations and, thus, the absolute value is demonstrated. The imaginary part of the gyration number corresponding to the rotation axis shows the strength of the “rotation” force in the dataset. The higher the imaginary part a dataset has, the stronger its structural rotations are. Typically (but not necessarily), a high imaginary part of a dataset’s gyration number is associated with proper structural rotations on the jPCA plot. B-D) The zoomed insets demonstrate the clear structural rotations pattern across datasets with different but high imaginary parts (above the diagonal line on the gyration plane as in condition 7). More “round” datasets are mostly concentrated near the rotation axis (inset B) while the more distorted ones (insets C, D) either have lower imaginary parts or significant real parts. E) The zoomed inset demonstrates datasets with low imaginary parts and high real parts of gyration numbers. The datasets demonstrate the deterioration of structural rotations as the imaginary part decreases culminating with the absence of rotational structure in the bottom part of the inset. F) The zoomed inset demonstrates datasets with small imaginary parts of the gyration number. The datasets demonstrate significant distortions of structural rotations reflecting the increase in the intrinsic dimensionality of data.

**Figure 5:**
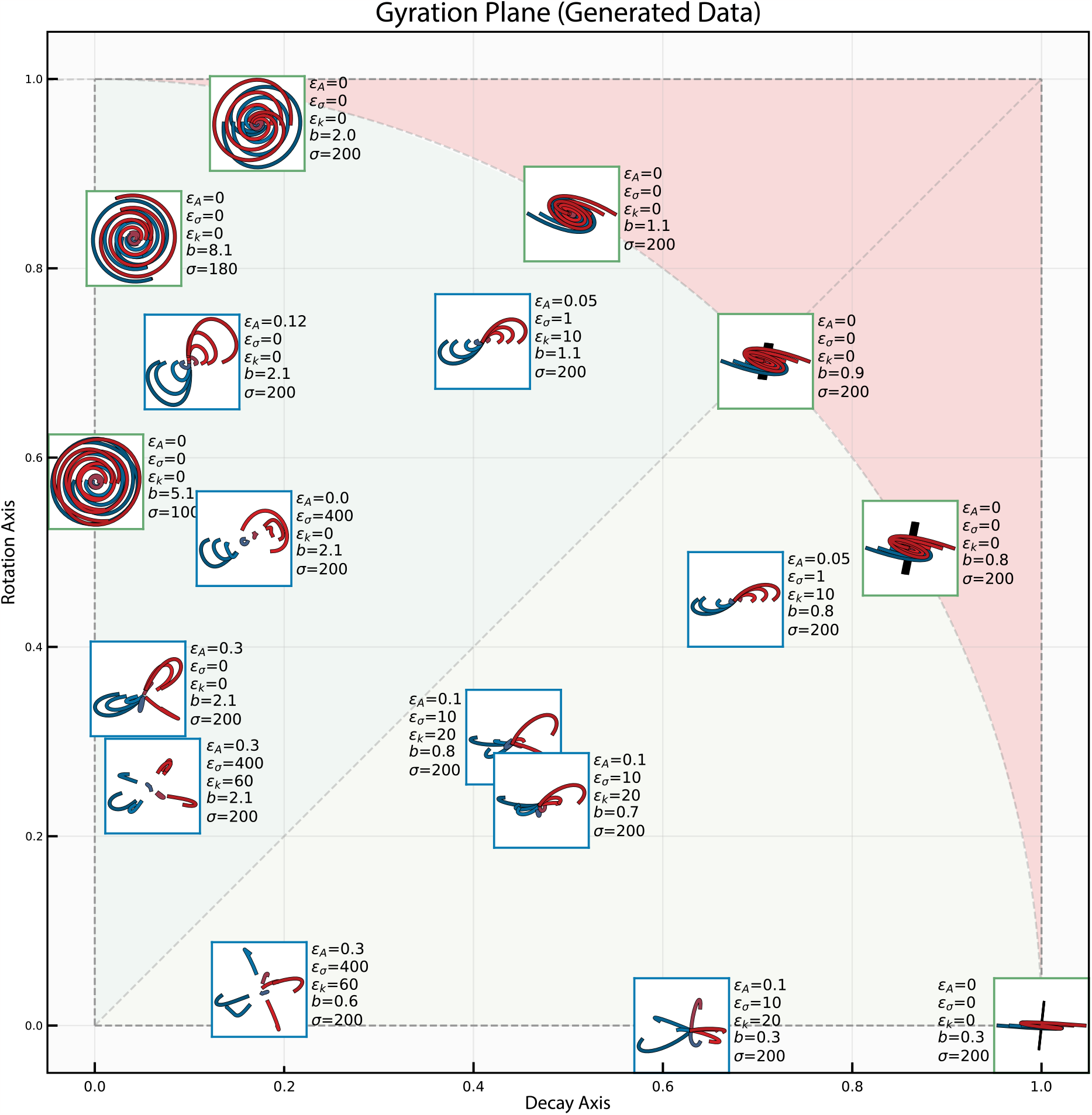
The gyration plane for simulated datasets. As in figure 4, insets demonstrate differently parameterized synthetic datasets including limit cases (located on the perimeter) of gyration number. Each inset is attributed with a corresponding set of parameters used for simulation (based on the eq. 9).

All datasets could be fit to the naive model of a travelling wave with a high *R*^2^ and the peri-event time histogram (PETH) for each dataset exhibited a travelling wave structure (Fig. S3). We also included three shuffled versions of the original dataset from [20], with the shuffling procedures implemented as described in the source article. In figures S4 and 4 the datasets where no rotational dynamics were reported in the original papers are delineated with red bounds. This include M1 and SMA recordings during grasping movements [22] and shuffled versions of dataset from [20].

The gyration plane can be empirically subdivided by the main diagonal into two areas. Datasets above the diagonal exhibit structural rotational dynamics, whereas those below it do not have structural rotational dynamics. However, this division is empirical and the rotational and decay axis represent more of a gradient than a strict transition from structural rotations to unstructured ones. Nevertheless, it is useful to note that the majority of datasets that are assumed to have rotational structure concentrate in the area above the diagonal.

Figure 4B-F shows zoomed gyration plane insets. The datasets in figure 4B exhibit prominent rotations with high score along the rotation axis. All the datasets in figure 4B are the neural recordings from M1 and PMd during hand-reaching movements.

The datasets in figure 4C have a higher score along the decay axis, but one can still observe prominent rotational dynamics visually. One of the main differences from the datasets in figure 4B is the diverse radius of trajectories. The conditions’ trajectories in each jPCA projection plot of figure 4C range from noisy conditions with small or non-rotational trajectory to conditions with a huge radius and clear rotational structure. Consistently with the gyration plane for the generated data (Fig. 5), the experimental datasets in figure 4C have more “stretched,” ellipsoidal shapes of rotations. Figure 4C contains data from somatosensory (S1, A2) and parietal (A5) areas and a recording of M1. All recordings were made during hand-reaching movements, as in figure 4B.

In figure 4D, various datasets from M1 and PMd during hand-reaching movements are shown, as well as two datasets recorded during hand-grasping tasks. All these datasets exhibit an average degree of structural rotations, with diverse radii of trajectories. The roundness of trajectories is also relatively small. These datasets are usually presumed to have a rotational structure.

Figures 4F and 4E contain the shuffled dataset of [20] (red-green color scheme) and PFC dataset (green-blue color scheme), which contain high frequency component, that perturbs structural rotational pattern. Thus, all datasets have a low rotational component in the data corresponding to a gyration number of approximately 0.15. Visually, we can evaluate those datasets as having weak and noisy structural rotational pattern.

All datasets in figure 4E are below the main diagonal. There are two shuffled datasets of [20], one from PFC, and the lowest is the neural activity of SMA recorded during grasping movements. All the datasets have been reported to have no rotational dynamics. One can see that the worsening of rotational dynamics (e.g. because of the shuffling procedures) is accompanied by a gradual decrease of the rotational component of the gyration number.

Overall, these results demonstrate the high similarity of datasets and the usefulness of the gyration number measure in characterizing structural rotational dynamics in neural data.

## 2. Discussion

Although rotational dynamics have been investigated for different datasets for over a decade, several issues remain unclear. Additionally, the two leading hypotheses, the dynamical model and the representational model, continue to coexist even though they offer very different interpretations. The former model posits that “the rotations of neural state space are an indicator of the dynamical system” [20], while the latter considers rotational dynamics as derivatives of representations [61, 70, 73].

Attempts to merge these two models have been made [51, 76, 77], framing the representational perspective as addressing “What motor parameters are involved?” and the dynamical system perspective as focusing on “How does neuronal activity evolve in time?” [11, 78].

The dynamical system approach faces the same challenges as the representational model: rotations exhibit diverse shapes and occur during various movement types, contexts, and behaviors (see Introduction), which are difficult to explain by either model or their combination [16, 17, 29, 51].

As noted in [70], interpreting rotational structure following dimensionality reduction should be conducted cautiously, as it is easy to misconstrue the true meaning of the data structure.

None of the interpretations named above can account for rotational phenomena from a data-driven perspective. Several approaches have been developed to study neural data rotations [79, 80], including the probabilistic jPCA method [81], which isolates neuronal rotations from motor cortex hypotheses.

The sequence-like response data model proposed by Lebedev et al. [61] comes closest to achieving this goal. While not a comprehensive model of how cortical dynamics develop, accounting for different phenomena (such as the high variability of single-neuron responses [55]), this model explains rotations by identifying a prominent structure in the data, namely, a travelling wave pattern.

In our study, we examined the idea of sequence-like responses further and expanded the analysis of rotational dynamics using a model-free perspective. We did not propose alternative models for generating neural activity or explanations for the underlying neurophysiological processes involved in voluntary movement generation. Instead, we demonstrated that a travelling wave is sufficient but not necessary for rotations to occur.

Furthermore, we proposed an explanation for the structure and properties in the data that give rise to rotational patterns. This explanation allows us to construct a natural complex-valued measure of rotations, a *gyration number*, that provides a strict definition of rotational dynamics. With this measure, we were also able to compare different real datasets of neural dynamics.

We distinguished between single-conditional rotations and structural rotations, and identified the necessary and sufficient requirements for the emergence of the former. We proposed that curvature, measured in high-dimensional space, is an advantageous measure of rotationess of the single trajectory. Strictly positive curvature indicates rotation in one direction, which is *necessary* for single-condition rotation. The absolute value of curvature points to particular twists of the trajectory and provides an intuition about the multidimensional behavior of the trajectory. However, a clear rotations of the original multidimensional single-condition trajectory could be distorted after dimensionality reduction methods (like PCA) due to improper projection (Figure 2B). The jPCA method better maintains single-condition rotation; however, like all projection techniques, it is prone to altering the actual curvature of trajectories when dealing with multiple conditions, as shown in Figure 2A.

While we acknowledge that low-dimensional manifolds can describe population dynamics, we emphasize that projective methods could generate false negatives and hide rotational dynamics in data where it actually exists.

Given that all datasets in this study exhibited travelling wave data patterns, we assumed that a simple travelling wave model can reproduce the rotational dynamics of single-condition data [61]. We demonstrated that such patterns imply a Toeplitz-like covariance matrix with oscillating eigenvectors (Fig. 3), and produce a clear horseshoe pattern in full coherence with PCA rotations of real neuron population data. Travelling wave data patterns also imply a skew-symmetric differential covariance matrix that yields an orthogonal rotational operator acting on the initial point of the data trajectory, as shown in the analytical solution of the jPCA problem (see eq. 6). Thus, we conclude that the travelling wave pattern is the *sufficient requirement* for a single-condition rotation.

Dealing with structural rotations, we wondered how different datasets with different numbers of conditions, neurons, and time points can be compared with each other in terms of the extent of their rotations. As an answer to this problem, we propose gyration number as a complex-valued measure of structural rotations in neural population datasets. This two-fold measure captures rotation and decay, with the imaginary part of the gyration number indicating the strength of rotations, and the real part measuring the stretchiness of trajectories in the dataset. The gyration number, visualized on a complex plane, enables model-agnostic comparison of different datasets in terms of their “rotationess” (Fig. 5), without requiring dimensionality reduction and preserving the full data information. Datasets recognized as rotating [20, 24, 25] exhibit a high imaginary part of the gyration number, while those with poor or no rotations have a high real part [22]. The gyration number provides a basis for meta-analysis of neural population data, although the rationale for such an analysis is subject to further discussion.

Upon the introduction of the measure of “rotationess”, we aimed to explore the possibility of different structural rotations. To achieve this, we generalized the travelling wave model for multiple conditions by introducing diversity in parameters typically observed in real data (see equation 9). Our observations revealed a crucial relationship between the intra-conditional noise (variance of the amplitudes of travelling waves within a condition) and inter-conditional noise (variance of the means of the conditions), which determines the existence of structural rotations (Supplementary figure S1). This relationship also affects the direction of the principal axes of a dataset and its corresponding jPCA (or PCA) representation. If the relationship favors the variance of the conditional means, a pronounced “sheaf” pattern appears on the jPCA plot before the cross-conditional average subtraction, and clear circular structural rotations emerge after the subtraction. Conversely, if the intra-conditional noise is high, the structural rotations vanish. It is also worth noting that the operation of subtracting the cross-conditional average serves only for the decoration of jPCA plot [61], not altering the dynamics of structural rotations but merely changes the starting point of about half of them to the opposite sign. Consequently, all conclusions regarding the ratio of the internal conditional noise and variance of the conditional means remain valid.

Furthermore, we found that data processing steps significantly affect rotational dynamics. The part of the signal used for projection can at least bias the visual evaluation of rotational consistency. Additionally, the relatively small number of conditions recorded in some studies, such as [45], can also affect rotational dynamics. The pre-processing step of applying a smoothing kernel to a spike train also greatly impacts rotational dynamics. For instance, we observed that slight changes in smoothing kernel width (from 10 to 20 ms) in the grasping movements dataset [22] resulted in either complete absence or presence of rotations for the neural activity recorded from the supplementary motor area (SMA) (not shown). All these peculiarities raise additional questions regarding the interpretation and comparison of different works on rotational dynamics.

We also demonstrated that structural rotations are not an artifact of the jPCA technique [61], but rather a property of the eigenstructure of the differential covariance matrix. To validate the importance of rotations, Churchland et al. and other researchers [18, 20, 45] employed various shuffling procedures. However, from a data-driven viewpoint, shuffling procedures are simply alternative ways of introducing additional intra-conditional noise to the initial dataset. Consequently, such procedures increase the dataset’s intrinsic dimensionality and modify the orientation of its principal axes, as we have described earlier (see Suppl. Fig. S1).

After considering the evidence presented, it is apparent that in many datasets, structural rotations are accompanied by a travelling wave pattern seen in the peri-event time histogram (PETH). The propagation speed, width, and consistency of this wave pattern provide a clear representation of the trajectories of conditions. Conversely, the absence of a travelling wave pattern results in the destruction of structural rotations. Supporting this, the naive travelling wave model can accurately describe real data, as demonstrated by its high *R*^2^ score and ability to reproduce structural rotations (Supplementary figure S3).

It is possible to hypothesize that in arm-reaching movements, the time order of neurons’ firing is fixed, while it is not the case for grasping movements. In datasets without rotational patterns, the behavior of each neuron varies significantly between conditions (as seen in the grasping dataset). This leads to different patterns of travelling wave propagation, resulting in the destruction of structural rotations in the low-dimensional projection. Thus, structural rotations in the data only indicate the propagating wave of activity in the neural population, providing no insight into the underlying dynamical system that generates motor cortex activity.

After examining rotational dynamics from a data-driven perspective, we have concluded that developing a model of cortical dynamics based on such limited data may be an endless process, and debates about the nature of structural rotations may be unproductive. This raises the important question: “Are the rotations of low-dimensional data projections encountered during numerous movements and in numerous brain areas useful?”

Although there have been attempts to use rotational dynamics for BCI and hand kinematic decoding [23, 82, 83], this approach has not been widely adopted.

It is important to keep in mind that the model of the rotations in the data is not a model of how the motor cortex generates volitional movements. The different goals that motivate model construction can affect model fitting and selection in dramatic ways [84]. For example, in [61], neural data from [20] was fitted to Lissajous curves with high precision. However, this does not imply that the underlying dynamics of the motor cortex are connected with those curves. Eventually, as Bassett et al. states [84], “Perhaps the first and most fundamental question that one can ask about a model is whether it is a simple representation of data or a theory of how the system behind the data might work.”

