## Supplementary Figures for "On the Rotational Structure in Neural Data"

Supplementary Materials for
**Unveiling rotational dynamics in neural data**
Ekaterina Kuzmina, Dmitrii Kriukov, Mikhail Lebedev

### Supplementary derivation A

In this supplementary derivation we demonstrate that the proposed running wave model 3 (see main text) generates skew-symmetric differential covariance matrix $\dot{X}^T X$, $X, \dot{X} \in \mathbb{R}^{t\times n}$. All derivations are conducted with *sympy* [85] package for python 3. Note that in real data cases we compute differential covariance matrix from finite dimensional $X$ and $\dot{X}$ that may cause edge-effects on corners of the matrix (see Fig. S5). To avoid the edge effects and generalize our considerations, instead of row finite vectors we consider row vector of time-dependent functions $X = [x_1(t), x_2(t), ...x_n(t)] \in \mathbb{R}^{1\times n}$ embedded in Hilbert space. For simplicity we show the case of $n = 3$ neurons, though all implications will hold for any finite $n$. Let $j = [0, ..., n-1] = [0, 1, 2]$ is an index of neuron then vector of functions. Substituting $j$ to the equation 4 (see main text) yields the following parametrization of $X$:

$$X = \begin{bmatrix} f(k(0), t, \sigma) & f(k(1), t, \sigma) & f(k(2), t, \sigma) \end{bmatrix} = \begin{bmatrix} e^{-\frac{(-a+t)^2}{\sigma^2}} & e^{-\frac{(-a-b+t)^2}{\sigma^2}} & e^{-\frac{(-a-2b+t)^2}{\sigma^2}} \end{bmatrix}. \tag{S1}$$

Parameters $a$, $b$, $\sigma$ are defined as in the equation 4. Next, we find time derivative vector $\dot{X} = \frac{d}{dt}X$ (note that all neurons are independent time-varying functions in our model and thus have diagonal Jacobian matrix):

$$\dot{X} = \begin{bmatrix} -\frac{(-2a+2t)e^{-\frac{(-a+t)^2}{\sigma^2}}}{\sigma^2} & -\frac{(-2a-2b+2t)e^{-\frac{(-a-b+t)^2}{\sigma^2}}}{\sigma^2} & -\frac{(-2a-4b+2t)e^{-\frac{(-a-2b+t)^2}{\sigma^2}}}{\sigma^2} \end{bmatrix}. \tag{S2}$$

Next, we need to find and analogue of differential covariance matrix for continuous time variable $t$. For any pair of neurons we define differential covariance as the scalar product of two time-varying functions, i. e. $\langle x_i(t), x_j(t)\rangle = \int_{-\infty}^{\infty} x_i(t)x_j(t)dt$. Using this definition the outer product of two vector of functions yields the square differential covariance matrix $\dot{X}^T X$.

$$\dot{X}^T X \to \langle \dot{X}, X\rangle = \begin{bmatrix} 0 & -\frac{\sqrt{2}\sqrt{\pi}be^{-\frac{b^2}{2\sigma^2}}}{2\sigma} & -\frac{\sqrt{2}\sqrt{\pi}be^{-\frac{2b^2}{\sigma^2}}}{\sigma} \\ \frac{\sqrt{2}\sqrt{\pi}be^{-\frac{b^2}{2\sigma^2}}}{2\sigma} & 0 & -\frac{\sqrt{2}\sqrt{\pi}be^{-\frac{b^2}{2\sigma^2}}}{2\sigma} \\ \frac{\sqrt{2}\sqrt{\pi}be^{-\frac{2b^2}{\sigma^2}}}{\sigma} & \frac{\sqrt{2}\sqrt{\pi}be^{-\frac{b^2}{2\sigma^2}}}{2\sigma} & 0 \end{bmatrix} \tag{S3}$$

One can see from the result for $n = 3$ that the obtained matrix is skew-symmetric ($X_{ij} = -X_{ji}$). We reproduced these derivations for other values of $n$ and obtained the same skew-symmetric pattern of matrix $\dot{X}^T X$. Interestingly, one can see that elements on the same offset diagonal of the matrix are the same that is the matrix is Toeplitz. We noted that this property holds only for even neuron activity functions like Gaussian in our model. What is more surprising, the elements of matrix do not depend on parameter $a$ which is in line with our numerical experiments. Thus, differential covariance and corresponding rotational dynamics does not depend on the initial wave shift (at least for our model assumptions).

### Supplementary figures

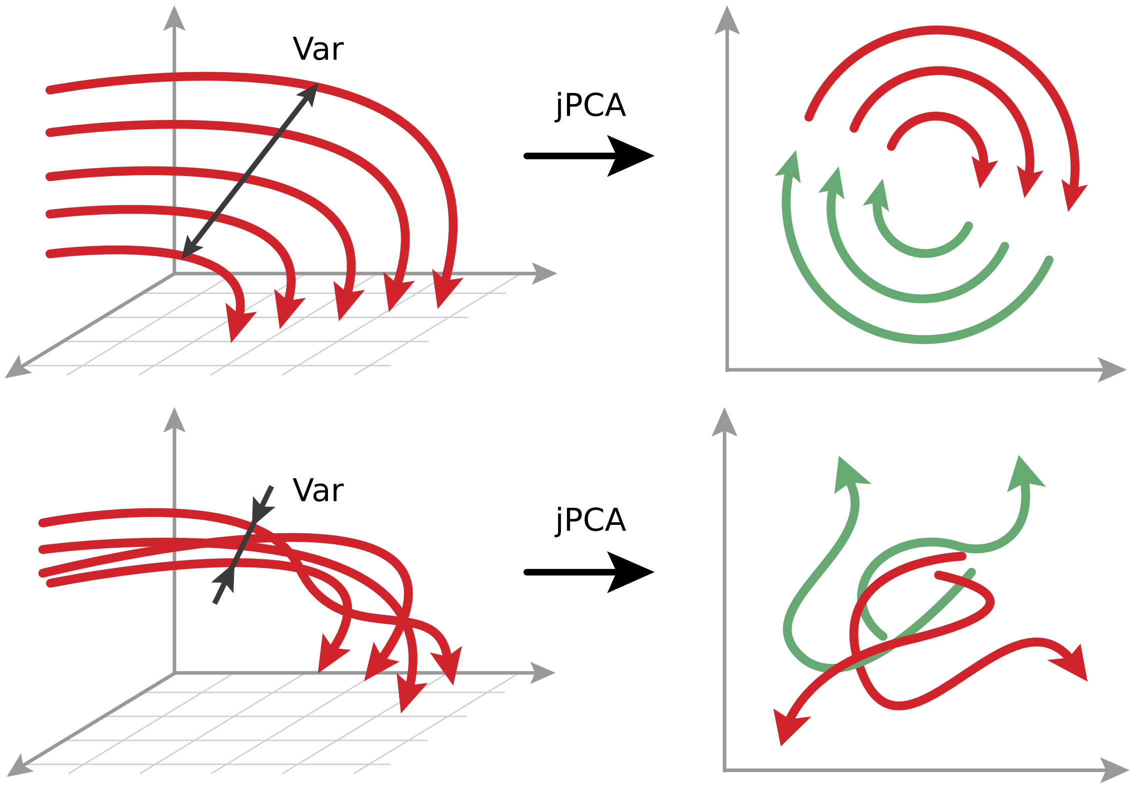


Figure S1: A sketch demonstrating the dependence of structural rotations occurrence of two factors: (i) high correlation between multidimensional conditions trajectories; (ii) sufficiently large variance between conditional-mean values (magnitude of trajectory coverage). Both factors are related to jPCA's property to catch a low-dimensional rotational manifold. The upper panel demonstrates the ideal case when the well-correlated conditions which have big enough variance maps to clear structural rotations. The bottom panel demonstrates the case when variance in condition means is not enough to form clear structural rotations in the jPCA plane.

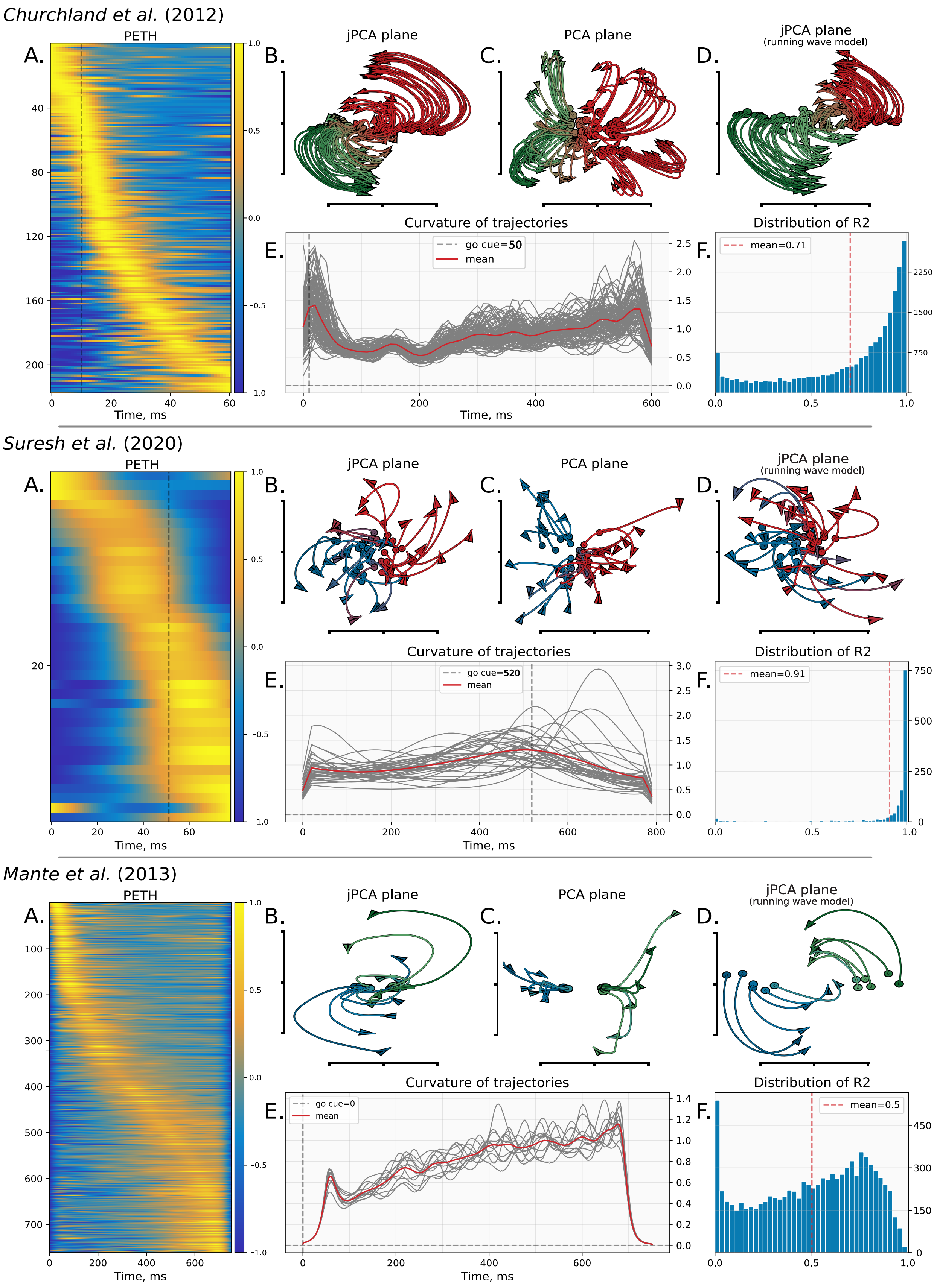


Figure S2: Datasets used in our research with some features. See details in the caption of figure S3

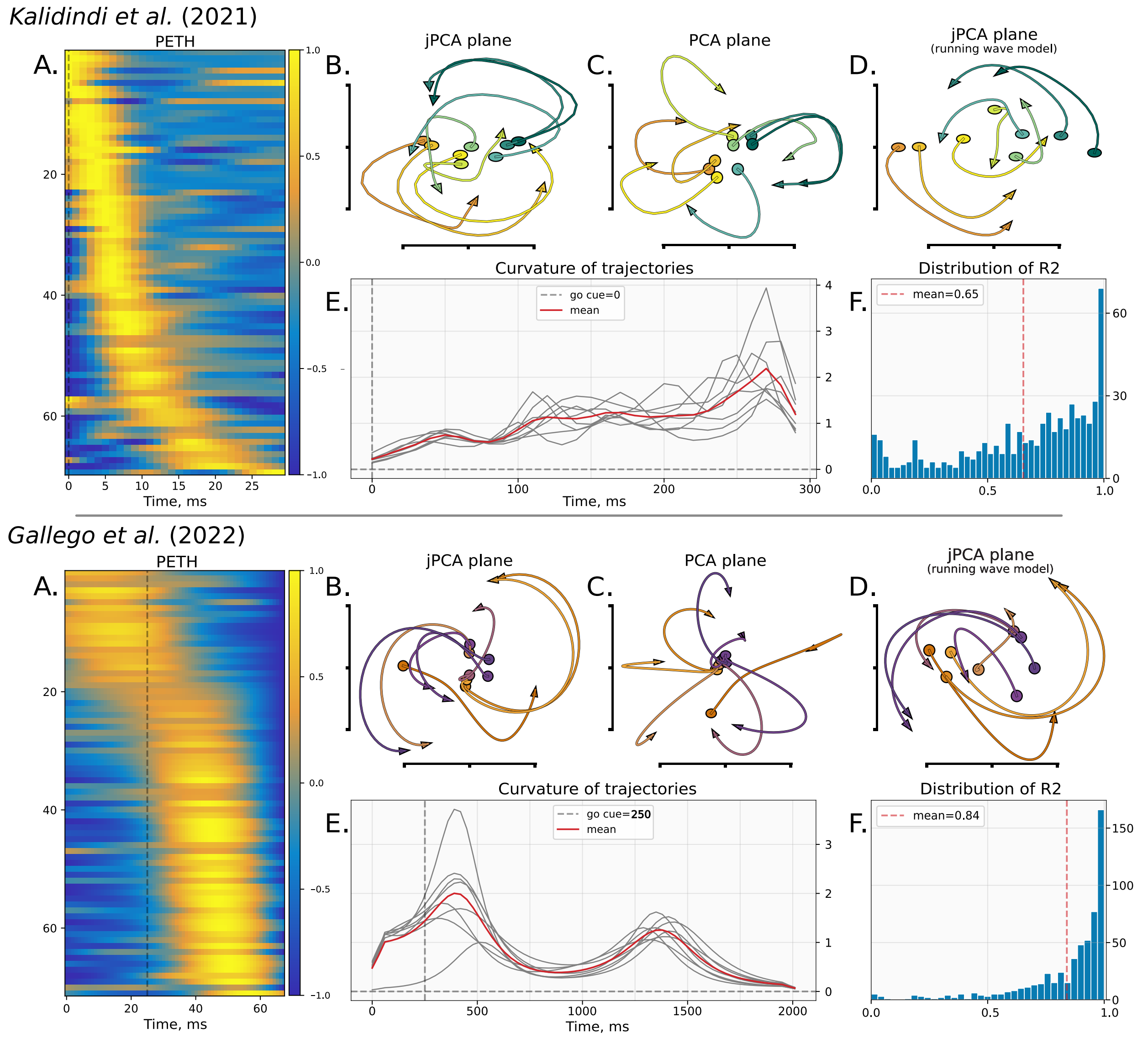


Figure S3: Datasets used in our research with some features. A) Peri-event time histogram of the dataset with the clear running-wave pattern. B) jPCA1-2 plot of the original dataset. C) PCA1-2 plot of the original dataset. D) jPCA1-2 plot of the running-wave fitted dataset. E) Curvature of each condition of the original dataset (grey lines) and average curvature (red line). F) $R^2$ histogram demonstrating a quality of running wave fit to neuron spike train data across all neurons and conditions. The Red dashed line shows the mean quality in terms of $R^2$.

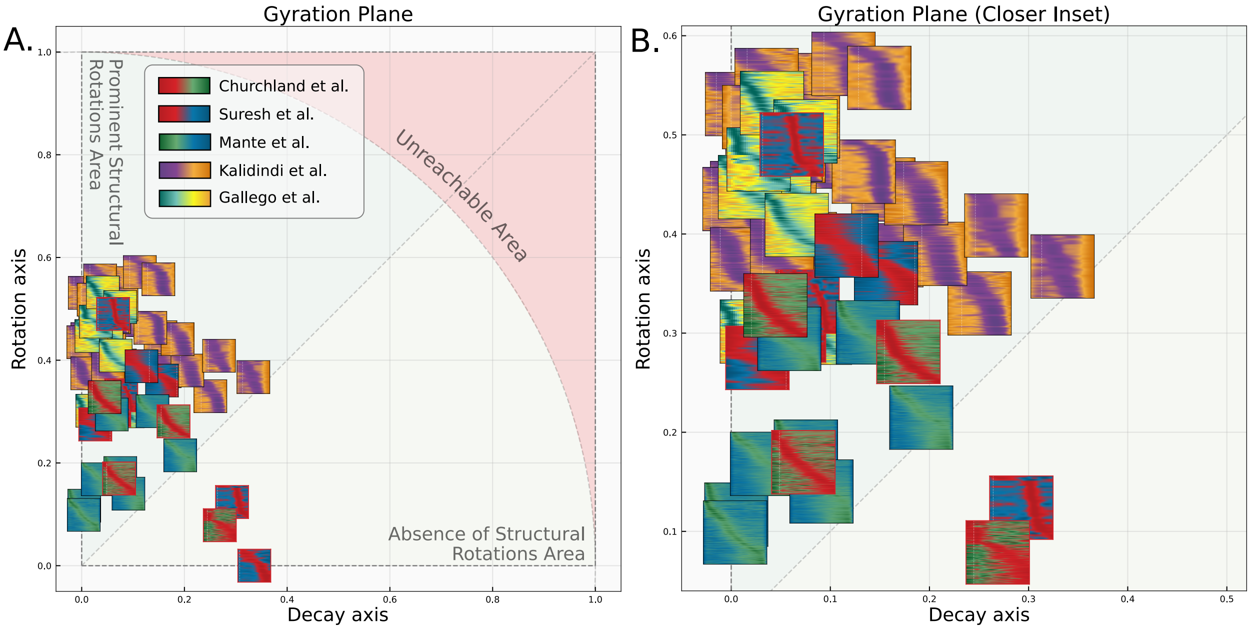


Figure S4: A. Gyration plane with PETH of datasets. The datasets location is the same as in gyration plane (Fig. 4). B. Zoomed inset of gyration plane with PETH.

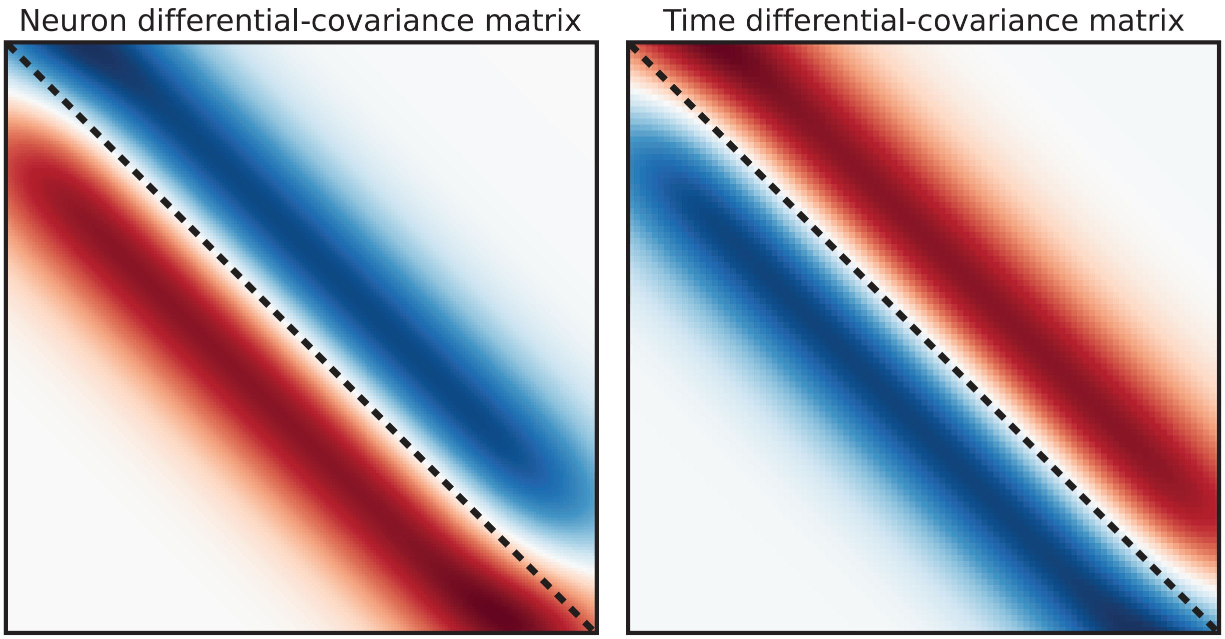


Figure S5: Differential covariance matrices for running wave data corresponding to the equation 3 (main text). Left, neuron differential covariance matrix $\dot{X}X^T$. Right, time differential covariance matrix $X^T\dot{X}$. Dashed gray line corresponds to the main diagonal of the matrix.

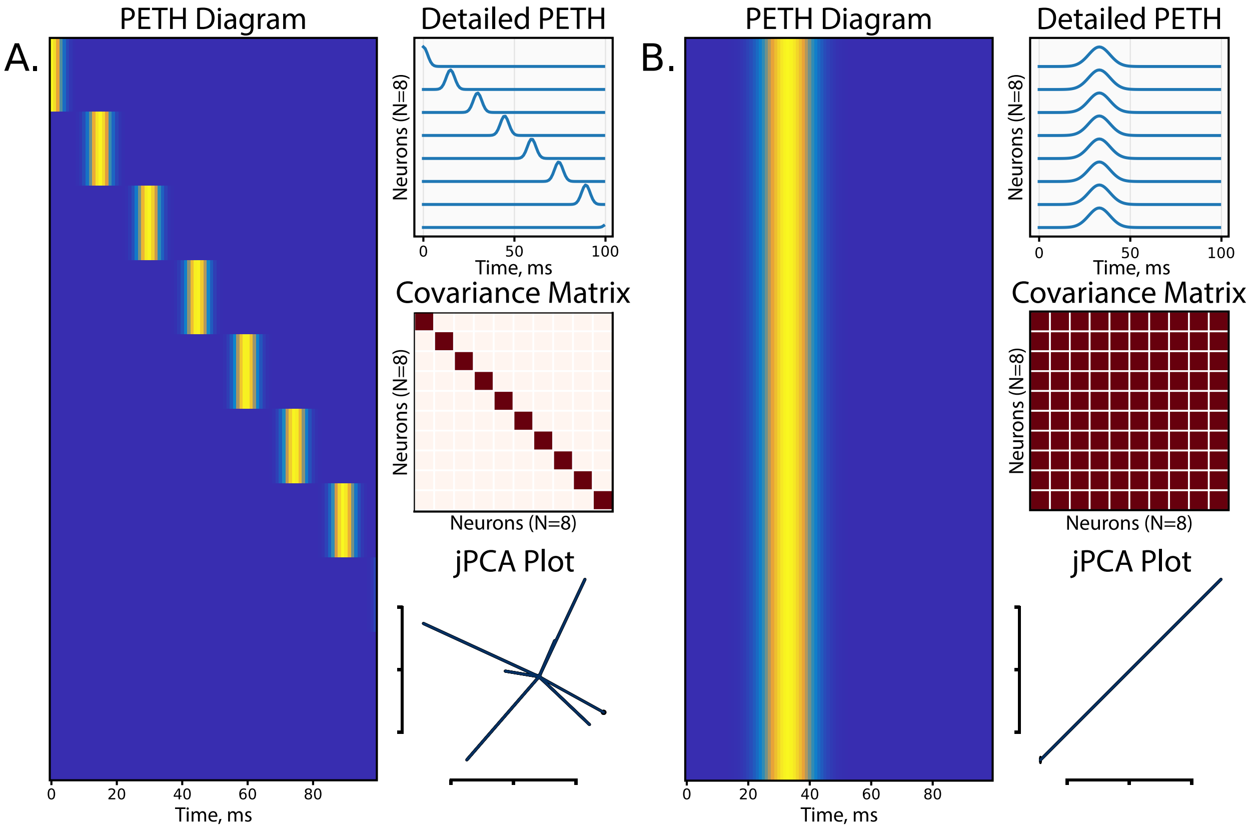


Figure S6: Limit cases of Gaussian running waves model. A) The case of uncorrelated neuronal spike-train waves is shown. Left: the corresponding PETH diagram in a form of a heatmap demonstrates disjoint waves of different neurons (high value of wave speed parameter). Upper right: another representation of PETH in a form of a time-series process. Middle right: the corresponding covariance matrix with diagonal structure demonstrates no correlation between neurons. Bottom right: jPCA dimensionality reduction technique maps spike train data to a set of orthogonal trajectories exhibiting no structural rotations. B) The case of fully correlated neuronal spike-train waves is shown. Left: the corresponding PETH diagram in a form of a heatmap demonstrates equal-phase waves (zero wave speed parameter) of different neurons. Upper right: another representation of PETH in a form of a time-series process. Middle right: the corresponding covariance matrix with unitary structure demonstrates the full correlation between neurons. Bottom right: jPCA dimensionality reduction technique maps spike train data to a set of collinear trajectories exhibiting no structural rotations.

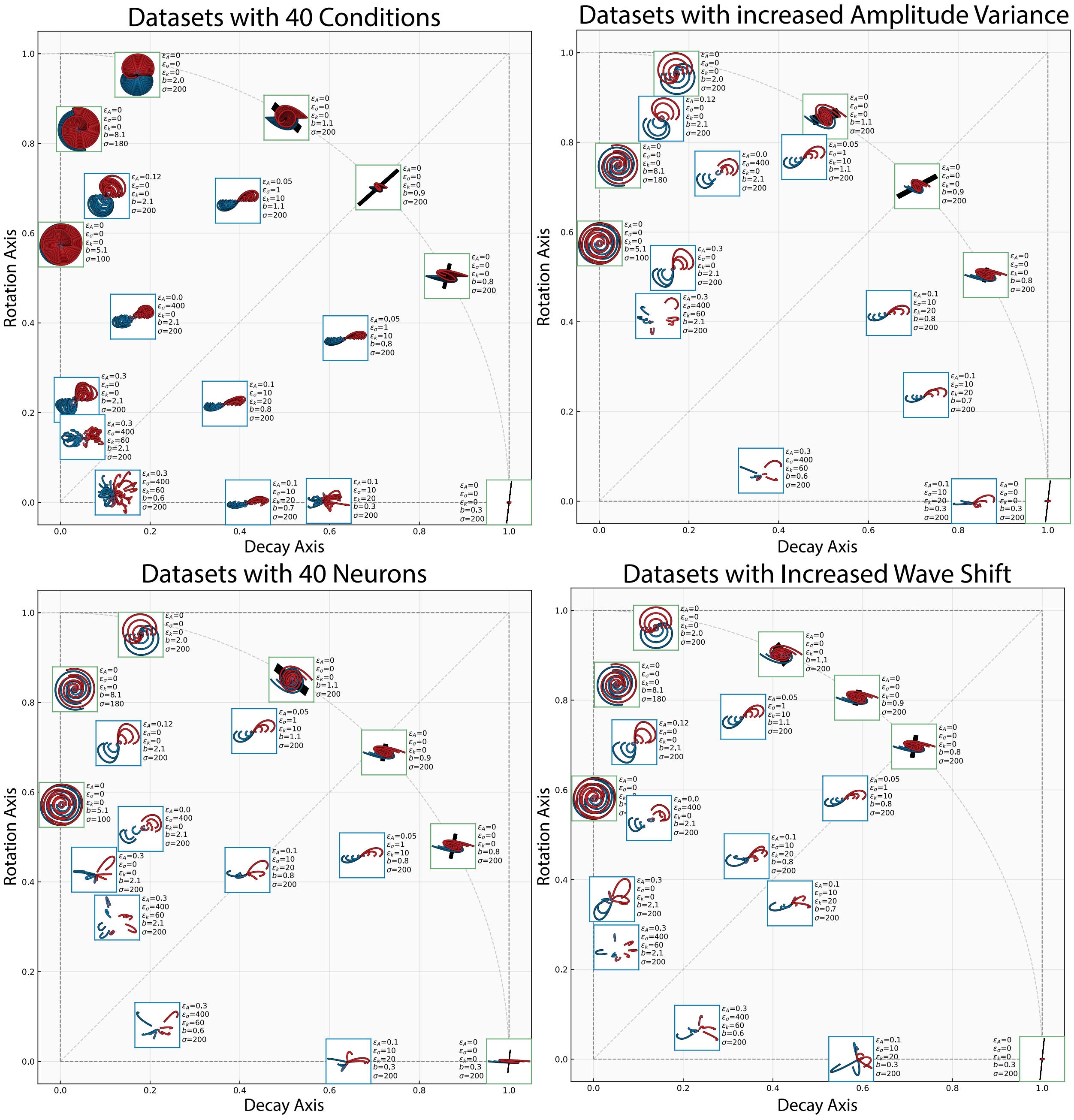


Figure S7: Figure S7: Gyration plane for synthetic datasets generated from different parameters. Condition perturbation parameters changed in an individual dataset are drawn near the corresponding dataset. Upper left: Datasets with 40 conditions in each dataset. Upper right: Datasets with increased amplitude variance $Var(A_l)$ (do not confuse with amplitude perturbation variance $\varepsilon_A$). Lower left: Datasets with 40 neurons in each condition of each dataset. Lower right: Datasets with increased initial wave shift $a$.
